# A comprehensive evaluation of ultraviolet irradiation at 254 nm and 222 nm in inactivating human noroviruses on surfaces

**DOI:** 10.64898/2025.12.18.695136

**Authors:** Hong Bai, Malcolm Turk Hsern Tan, Jiangyong Hu, Walter Randazzo, Dan Li

## Abstract

Human norovirus (hNoV) presents significant public health challenges due to its low infectious dose and environmental persistence. This study compared the inactivation efficacy of ultraviolet C irradiation at 254 nm (UV 254) and far-UVC radiation at 222 nm (UV 222) against four hNoV strains and two surrogate viruses, Tulane virus (TV) and bacteriophage MS2. A symptom scoring assay was developed to assess hNoV infectivity following microinjection into zebrafish embryos, being used in combination with reverse transcriptase quantitative PCR (RT-qPCR), long-range RT-qPCR, and RNase-treated RT-qPCR. With a general laboratory set-up of viruses being suspended in deionized water droplets in Petri dish, UV 222 was demonstrated with comparable, if not superior, performance in reducing hNoV infectivity and RNA integrity, and was significantly more effective than UV 254 in damaging viral capsids. MS2 exhibited inactivation patterns similar to hNoVs, whereas TV was markedly more resistant to UV 222. The performance of UV 222 was consistent in inactivating hydrated viruses on both stainless steel and porcine skin surfaces. However, the efficacy of UV-222 was substantially more reduced when virus inocula were dried or mixed with simulated vomitus containing high levels of organic matter, compared with UV-254. No evidence of viral adaptation or persistent genomic diversification was detected following repeated sublethal UV exposures. Taken together, UV 222 can be regarded as a promising technology in surface disinfection, especially for hNoV control, meantime keeping safe for human exposure.

**Importance:** Human norovirus, the main cause of foodborne illness and non-bacterial gastroenteritis, can be transmitted through human-to-human contact. Indirectly, food or food related surfaces are readily contaminated by hNoV, completing the transmission route. So far, no standard cultivation tool is available for detecting viable hNoV, resulting in the challenges of evaluating inactivation effectiveness of various disinfection technologies, including UV 222 treatments. The significance of our study lays in attempts to quantify hNoV infectivity loss of four strains using zebrafish model during UV 222 and 254 treatments, together with the underlying antiviral mechanisms indicated by three different types of reverse transcription qPCR methods. In addition, the concerns over possible emergence of variant were subdued by genome wide sequencing results after consecutive UV exposures and passaging *in vivo* zebrafish model.

**Synopsis:** UV 222 is recommended to be applied after surface cleaning and ideally on moist surfaces.

## Introduction

Human norovirus (hNoV), a non-zoonotic pathogen, has become the leading cause of foodborne illness and non-bacterial gastroenteritis globally over the past decade (1). While infection are typically self-limiting in healthy adults, hNoV can lead to severe and prolonged illness in immunocompromised individuals, such as young children or elderly people (2). As a member of the *Caliciviridae* family, hNoV possesses a single-stranded, positive-sense, polyadenylated (poly-A) RNA genome of approximately 7.5 kilobases, comprising three open reading frames (ORFs). ORF1 encodes the six nonstructural proteins, while ORF2 and ORF3 respectively encode the major (VP1) and minor (VP2) capsid proteins (3).

HNoV transmission occurs primarily through human-to-human contact either directly or indirectly via contaminated food, water, or environmental surfaces. Being highly contagious, hNoV can spread rapidly in closed or semi-enclosed environments such as cruise ships, leading to large-scale outbreaks. For example, in 2025, the U.S. Centers for Disease Control and Prevention (CDC) has reported 19 hNoV outbreaks on cruise ships (4). Due to its exceptional environmental stability, eliminating hNoV contamination once it occurs is extremely challenging. Ultraviolet C (UVC) irradiation at 254 nm (UV 254) has been extensively used for its germicidal properties in surface disinfection (5); however, its application is limited to unoccupied environments due to potential harm to human skin and eyes. Recent advances in far-UVC technology, particularly krypton-chloride (KrCl) excimer lamps emitting at 222 nm (UV 222), present a promising alternative. Far-UVC at 222 nm has demonstrated effective antimicrobial activity against various pathogens (6, 7), while keeping safe for human exposure as a result of limited penetration (8). These unique properties open avenues for continuous surface disinfection in occupied spaces, and hand disinfection (9, 10).

For a long time until very recently, due to the lack of a reliable cell culture system for hNoVs, a series of cultivable surrogates including murine norovirus (MNV), Tulane virus (TV), bacteriophage MS2 and feline calicivirus (FCV) have been used to evaluate hNoV inactivation (11–13). However, the suitability using these surrogates to accurately reflect hNoV behavior remains questionable (14). The susceptibilities of different viruses, including those among various hNoV genotypes, have been shown to vary considerably under different inactivation treatments, such as heat, high hydrostatic pressure, and exposure to disinfectants like chlorine, ethanol, and hydrogen peroxide (11, 12, 15). Previously, UV 222 has been reported to effectively reduce the infectivity of MNV and FCV to levels comparable to those achieved with UV 254 (16, 17), and has shown superior inactivation efficacy against MS2 compared to UV 254 (18). However, the direct antiviral effect of UV 222, its comparative efficacy relative to UV 254, and the underlying inactivation mechanisms have yet to be investigated for genuine hNoV strains (7).

Several years ago, a notable breakthrough came with the successful cultivation of hNoV with human intestinal enteroids (HIE) model. Being labor-intensive, expensive and requiring human biopsy specimens (13, 14), the use of the HIE model in hNoV inactivation studies has been mainly limited to validation of “complete inactivation” so far without generating quantitative data in virus titers or log reduction values. (1, 19). More recently, zebrafish larvae gained attention as a novel model, as they were shown to support hNoV replication following microinjection into their yolk sac (20). Building upon this advancement, our group has demonstrated that zebrafish embryos offer an improved efficiency and robustness for hNoV propagation (21). This model provides a more practical and accessible platform for evaluating the efficacy of inactivation strategies in a quantitative way (21, 22).

In this study, we established a new approach evaluating hNoV infectivity based on the zebrafish post-infection symptom scoring, with further improved efficiency compared with the past approaches based on virus loads (21, 22). With this new method, we investigated how UV irradiations at 222 and 254 nm inactivate hNoV strains GII.2[P16], GII.3[P12], GII.4 Sydney[P16] and GII.17[P31]. In the meantime, two surrogates TV and MS2 were tested in parallel. In addition, the capsid and genomic integrity of the viruses before and after treatment were evaluated by reverse transcriptase quantitative polymerase chain reaction (RT-qPCR), long range RT-qPCR (LR-RT-qPCR) and RNase RT-qPCR (RNase-RT-qPCR), shedding light on inactivation mechanisms.

Subsequently, to assess the practical applicability of UV 222 for hNoV inactivation on surfaces, its inactivation efficiency was examined on stainless steel, a representative operational surface commonly found in food processing environments, under both hydrated and dried conditions. HNoV-infected food handlers, as well as asymptomatic carriers, play a crucial role in the transmission of hNoVs, underscoring the importance of proper hand hygiene, as demonstrated by numerous studies employing various experimental approaches (23, 24). Hand sanitization represents an effective intervention for controlling viral transmission, particularly under conditions where conventional handwashing is impractical, such as during produce harvesting in the absence of potable water. Since UV 222 has been demonstrated to be safe for skin (25, 26), we further evaluated viral inactivation on porcine ear skin, a commonly used surrogate model for human skin (27). Since the vomit-oral transmission route represents a key pathway for hNoV spread, the influence of simulated vomitus matrix on UV inactivation efficiency was also investigated to better reflect real-world contamination scenarios.

Last, hNoV, like other RNA viruses, is characterized by high mutation rates, largely resulting from the low fidelity of RNA polymerases and the absence of proofreading mechanisms during replication (28). Previous studies on MNV showed that repeated sublethal exposure to disinfectants can drive the emergence of resistant variants with specific mutations, suggesting the potential for adaptive responses in hNoV under environmental pressures (29, 30). As UVC irradiation, particularly at 254 nm, has been extensively reported to induce primary damage to nucleic acids through the formation of specific photoproducts such as cyclobutane pyrimidine dimers and pyrimidine-(6–4)-pyrimidone photoproducts (31), we analyzed in this study the mutation rates of hNoV sublethally treated with UV irradiation at 222 and 254 nm during serial passaging in zebrafish using next-generation sequencing (NGS), thereby assessing the long-term impact of UV irradiation on viral evolution.\

## Results

### Establishment of symptom scoring method to evaluate hNoV infectivity

Following the protocol established by J. Van Dycke et al. (20), our previous studies assessing hNoV viability in response to inactivation treatments were based on viral load measurements. Specifically, hNoV genome copies were quantified in pooled samples of ten zebrafish larvae at 3 dpi and compared to the genome copies initially injected into each group at 0 dpi. A sample was considered positive if the viral genome copies increased by more than 2 logs within three days (21). Although this tool achieved to evaluate the efficacy of hNoV inactivation quantitatively (21, 22), this method failed to reveal a clear dose-response. We believe this is attributable to substantial individual variability in viral loads among zebrafish following infection, as observed in our recent study (data not shown). Moreover, the method entails a highly labor-intensive workflow, including virus harvesting, RNA extraction and RT-qPCR quantification, together with considerable costs of reagents.

In this study, we explored the potential of assessing virus viability through symptom scoring in zebrafish post-infection. This approach was inspired by the TCID₅₀ assay, a classical method for titrating cell-cultivable viruses which determines the viral concentration at which 50% of infected cells exhibit CPE, rather than relying on viral load measurements. As shown in Fig. 1, at 3 dpi, the symptoms of infected zebrafish larvae were categorized into four distinct scoring groups based on morphological and behavioral observations under a stereo microscope. We evaluated the scoring system over four different hNoV strains (GII.2[P16], GII.3[P12], GII.4 Sydney[P16] and GII.17[P31]) in three 10× serial-diluted concentrations. As a result, although some of the zebrafish might have developed asymptomatic infection, the three biological replicates demonstrated excellent linearity and repeatability for the sum of symptomatic scores from eight zebrafish larvae, with correlation coefficients between 0.82 to 0.94 from the regression lines (Fig. 2).

**Fig. 1.**
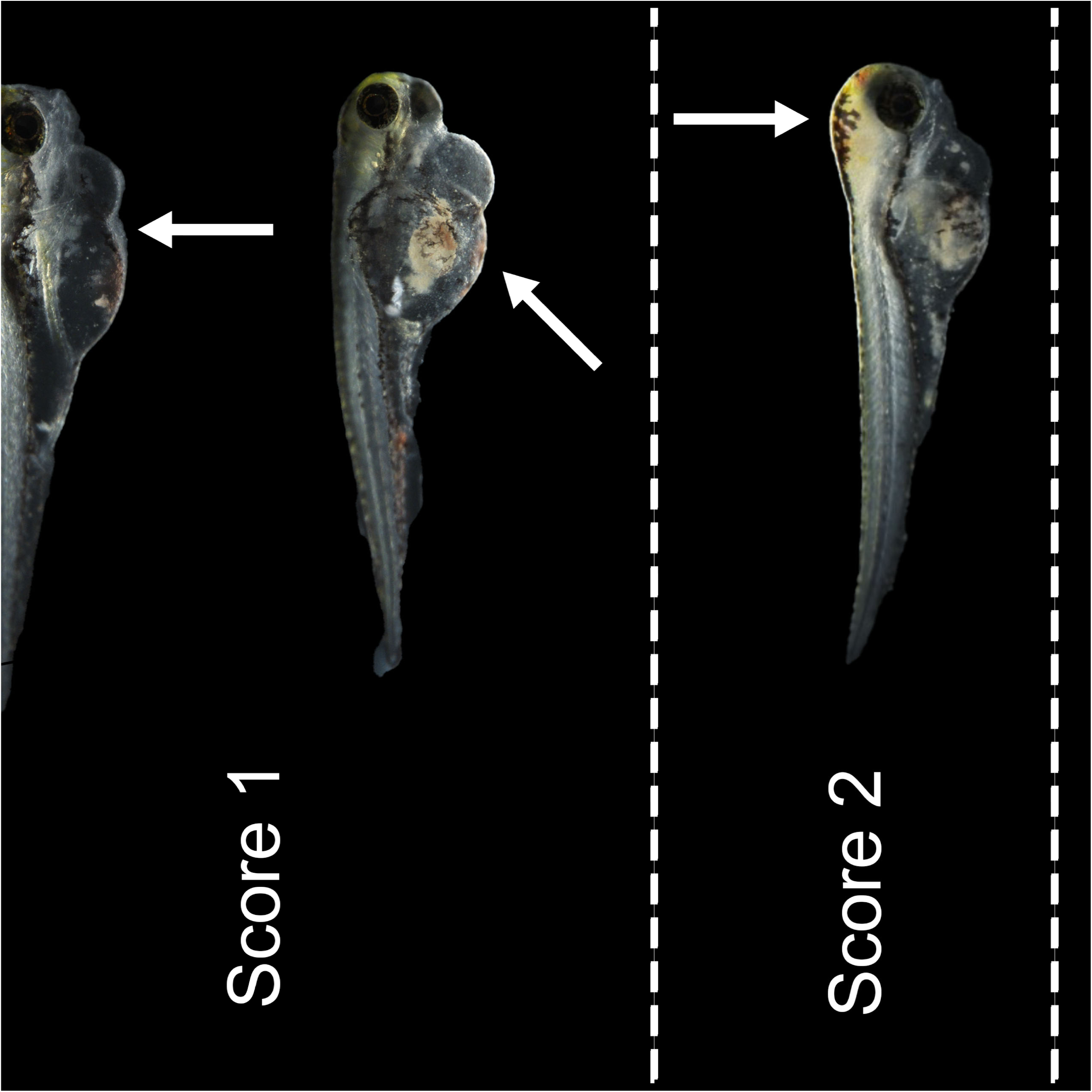
The morphological and behavioral classification of infected zebrafish larvae at 3 dpi for scoring. The standards for classifying were as following: 0-Normal fish: clear head and yolk sac area, and no sign of edema; 1-Mild symptoms: pronounced edema in the yolk sac or pericardial region, reduced responsiveness, or pale staining or opacity occurs in yolk sac; 2-Severe symptoms: pale staining or opacity occurs in head and yolk sac, pronounced edema in the yolk sac or pericardial region, lack of responsiveness; 3-Death and deformation: complete tissue degradation or loss of structural integrity.

**Fig. 2.**
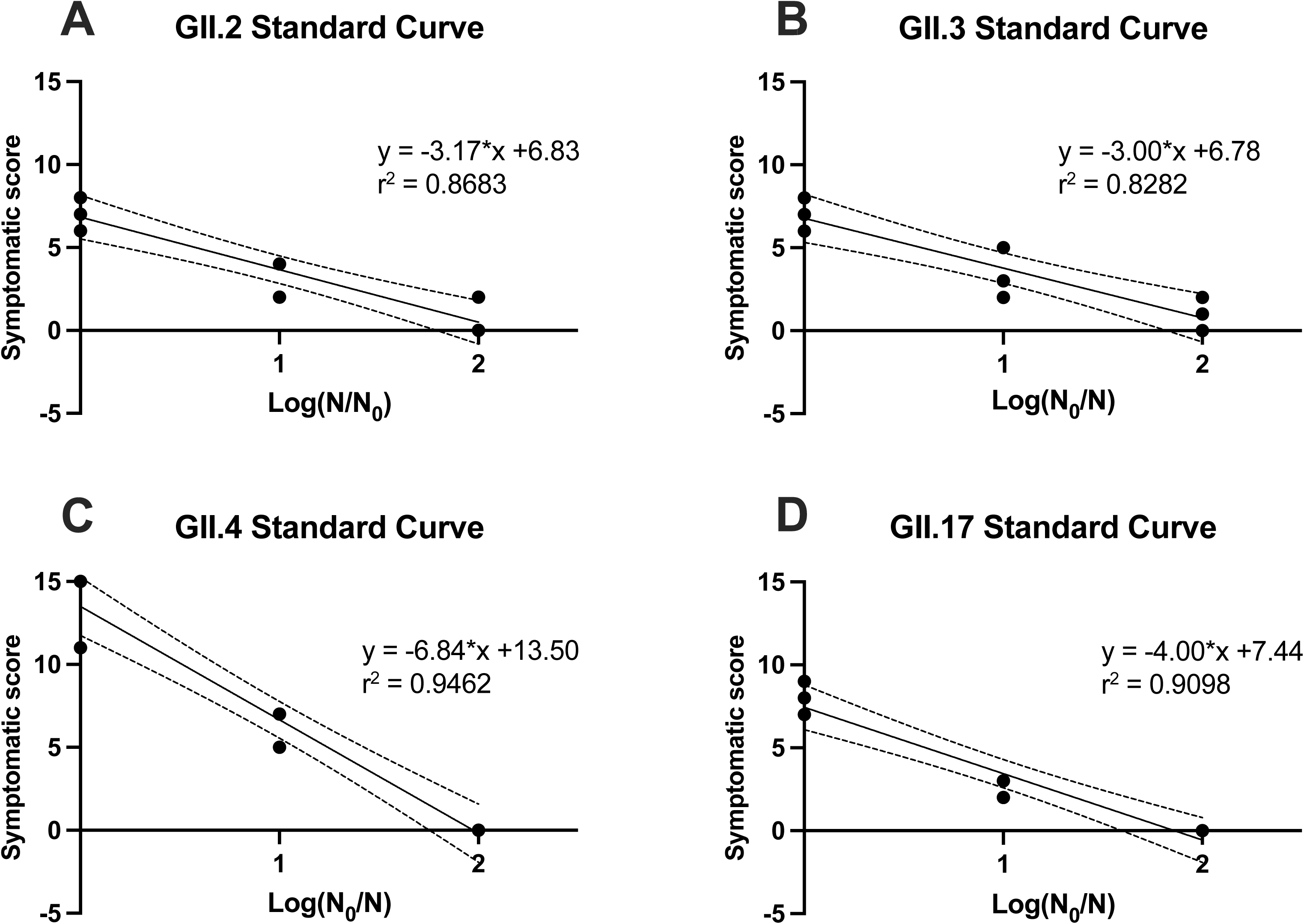
The single linear regression of zebrafish larvae symptom scores of eight larvae and log reductions for hNoV GII.2[P16], GII.3[P12], GII.4 Sydney[P16] and GII.17[P31] in biological triplicates.

### UV inactivation of hNoVs and surrogates

Based on the infectivity assay established as described above, the inactivation of four hNoV strains (GII.2[P16], GII.3[P12], GII.4 Sydney[P16] and GII.17[P31]) by UV 222 and UV 254 at 7 mJ/cm^2^ and 70 mJ/cm^2^, respectively, was evaluated. At 7 mJ/cm^2^, UV 222 exhibited slightly higher efficacy in inactivating hNoV GII.2[P16] (> 2.16 log reduction) than UV 254 (1.84 ± 0.32 log reduction), whereas no significant difference was observed between the efficacies of UV 222 and UV 254 on hNoVs GII.3[P12], GII.4 Sydney[P16] or GII.17[P31] (*p* > 0.05) (Fig. 3A). UV 222 and UV 254 at 70 mJ/cm^2^ both induced higher inactivation of hNoVs than 7 mJ/cm^2^, reaching the limits for all four strains (>2.16, >2.26, >1.98, >1.86 log reductions for GII.2[P16], GII.3[P12], GII.4 Sydney[P16] and GII.17[P31] respectively) (Fig. 3B). As expected, RT-qPCR, targeting on a short amplicon of the virus genome, reflected minimal reductions by the inactivation treatment (Fig. 3C, 3D). Although amplification of full-length genomic RNA by qPCR suffers from the limitation of insensitivity, separating the PCR amplification site and RT priming site within the virus genome stood as an effective alternative to measure the virus genome integrity (32, 33). In this study, the LR-RT-qPCR, quantifying the longer length of cDNA from intact RNA, reflected much clearer dose response of the UV treatment on the virus genome integrity (Fig. 3E, 3F). The influence of UV 222 and UV 254 on hNoV genomes were comparable for all of the tested scenarios (*p* > 0.05; Figure 3E, 3F) except for hNoV GII.3[P12], UV 254 at 7 mJ/cm^2^ induced significantly higher reduction than UV 222 at 7 mJ/cm^2^ (*p* < 0.05; Fig. 3E).

**Fig. 3.**
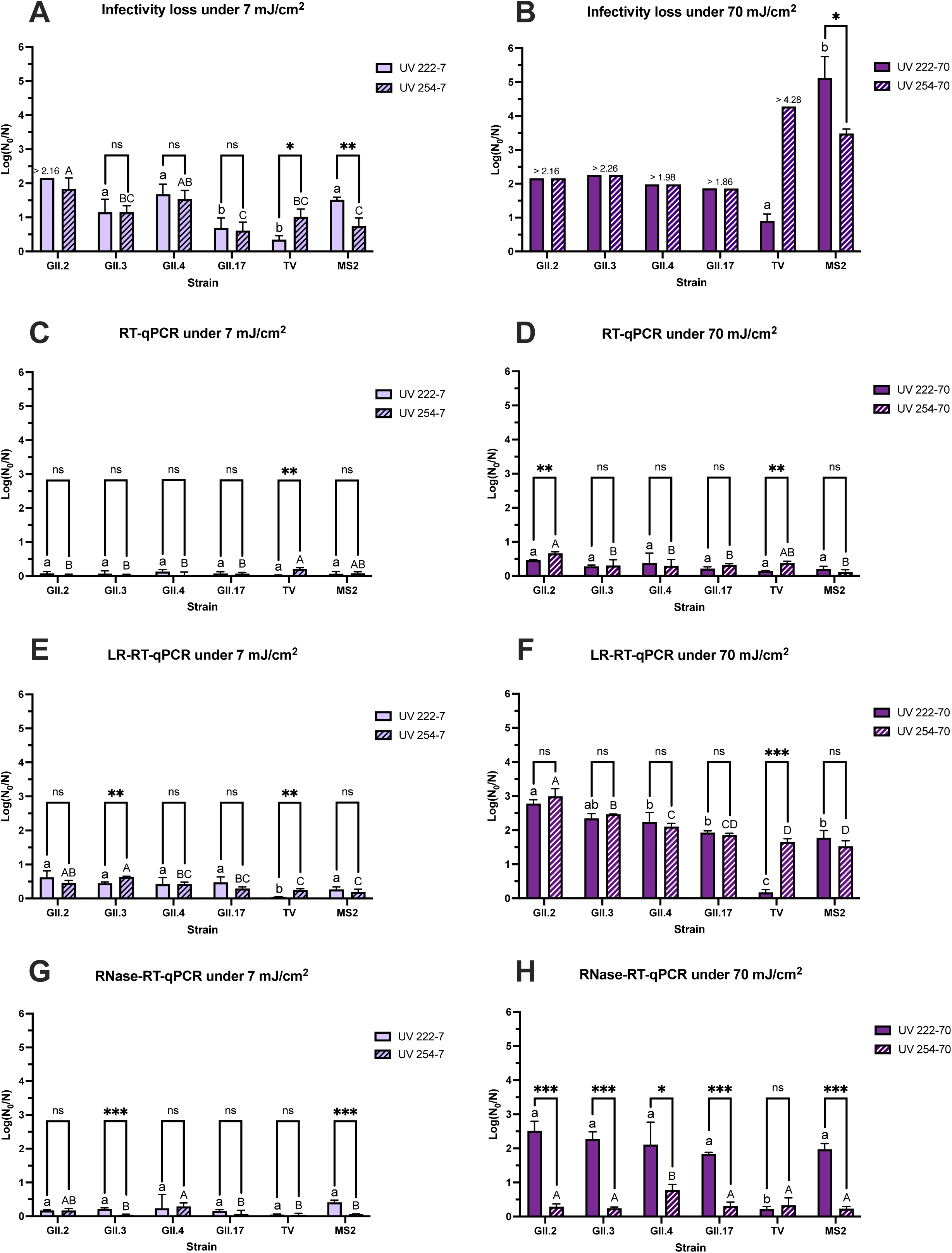
Under UV 222 and UV 254 treatment at 7 (A, C, E, G) and 70 mJ/cm^2^ (B, D, F, H), the log reductions of hNoV strains (GII.2[P16], GII.3[P12], GII.4 Sydney[P16] and GII.17[P31]) and surrogates (TV and MS2) through infectivity assays, RT-qPCR, LR-RT-qPCR and RNase-RT-qPCR. For pairwise comparison of UV treatment within each strain, differences are expressed as * (*p* < 0.05), ** (*p* < 0.01), *** (*p* < 0.001), ns (not significant). The letters above each column represents the differences among each UV treatment group, lowercase letters for UV 222, capital letters for UV 254. The log reductions with different letters suggest statistically significant differences (*p* < 0.05), while those with the same letters indicate not significantly different (*p* > 0.05).

The RNase-RT-qPCR method was established and tested on multiple enteric viruses to reflect the virus capsid integrity and predict the viral infectivity, based on the assumption that the free RNA and RNA in broken virus capsids which are exposed to RNase digestion would no longer be infectious (34, 35). In this study, RNase-RT-qPCR detects RNA in an intact viral capsid, indicating the integrity of virus capsid after UV treatments by digesting the un-protected RNA prior to RT-qPCR detection. Very interestingly, UV 222 demonstrated remarkably higher efficacies in damaging hNoV capsid than UV 254 (Fig. 3G, 3H). Significantly higher reductions were detected within all four tested hNoV strains by UV 222 than UV 254 at 70 mJ/cm^2^ (*p* < 0.05; Fig. 3H).

The four tested hNoV strains indeed showed some variabilities toward the UV inactivation, with GII.2[P16] being the most susceptible strain and GII.17[P31] the most persistent strain against both UV 222 and UV 254 as reflected by multiple assays (Fig. 3). However, all four hNoV strains showed consistent trends regarding the comparisons between the efficacies and mechanism of UV 222 and UV 254, supporting a clear conclusion that UV 222 has comparable efficacy (if not better) in inactivating hNoVs and is much more efficient in damaging hNoV capsid protein than UV 254. As for the surrogates, MS2 closely resembled the responses of genuine human NoV strains as measured by all four different methods (Fig. 3), whereas TV showed distinctively opposite reactions against the UV treatment in comparison with the rest of the tested viruses (Fig. 3). More specifically, at the same doses, UV 254 was much more efficient in inactivating TV than UV 222 (Fig. 3A, 3B). As indicated by the LR-RT-qPCR results, UV 254 and UV 222 were equally effective in damaging the genomic materials of hNoVs and MS2 (*p* > 0.05), whereas only UV 254 but not UV 222 was able to degrade the RNA of TV (*p* < 0.05; Fig. 3E, 3F). UV 222 was very efficient in damaging the capsid protein of hNoVs and MS2 in comparison with UV 254 (*p* < 0.05), but UV 222 and UV 254 were equally ineffective on the capsid protein of TV as suggested by the RNase-RT-qPCR results (*p* > 0.05; Fig. 3H).

### Application of UV treatments in surface disinfection

HNoV GII.17[P31] was selected for the surface disinfection evaluation as it was the most UV-resistant strain among those tested against UV 222 and UV 254, based on the comprehensive assessment shown in Fig. 3, thereby representing the worst-case scenario. LR-RT-qPCR was employed as it best reflected changes in hNoV infectivity among the three molecular methods tested, as shown in Fig. 3, while also capturing a broader range of reductions than the infectivity assay. MS2, which displayed resembled responses of genuine human NoV strains against UV 222 and UV 254 as shown in Fig. 3, was measured in parallel with infectivity-based plaque assay.

As shown in Fig. 4, when the virus inocula were treated in a hydrated form, UV 222 and UV 254 achieved comparable viral reductions on both stainless steel and porcine ear skin (*p* > 0.05). These reductions were also consistent with those observed in Petri dishes, as shown in Fig. 3. In contrast, when the virus inocula were dried on the surfaces prior to treatment, the efficacy of both UV 222 and UV 254 was markedly reduced. UV 254 exhibited significantly greater inactivation efficiency than UV 222 against hNoV GII.17[P31] on stainless steel and MS2 on both surfaces when the viruses were dry (*p* < 0.05; Fig. 4). Likewise, the presence of simulated vomitus compromised the performance of both UV 222 and UV 254. UV 254 exhibited significantly greater inactivation efficiency than UV 222 against both hNoV GII.17[P31] and MS2 on both surfaces when the viruses were mixed with simulated vomitus with high organic loads (*p* < 0.05; Fig. 4).

**Fig. 4.**
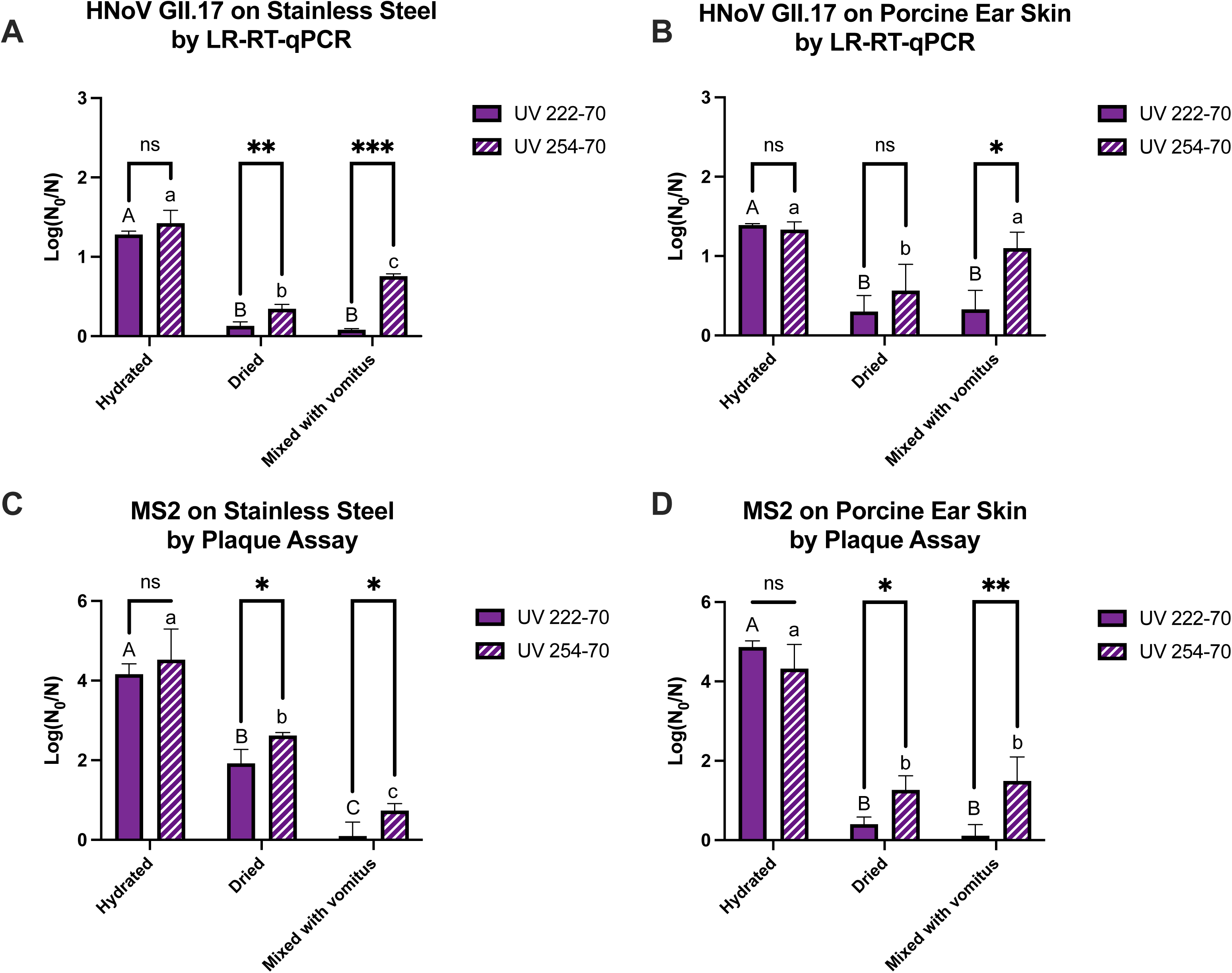
Under UV 222 and UV 254 treatments at 70 mJ/cm^2^, the log reductions of hNoV GII.17[P31] LR-RT-qPCR results and MS2 infectivity in hydrated or dried forms, as well as in hydrated vomitus on stainless steel surfaces (A and C) and on porcine ear skin (B and D). For pairwise comparison of UV treatment within each treatment and each state of virus, differences are expressed as * (*p* < 0.05), ** (*p* < 0.01), *** (*p* < 0.001), ns (not significant).

### Evaluation of hNoV adaptation and mutation after repeated UV treatments

Based on the LR-RT-qPCR results presented in Fig. 3 and 4, both UV 222 and UV 254 treatments affected the integrity of the hNoV genome. To unveil the effect of UV lights on genomic mutation, two aliquots of GII.4 Sydney[P16] clinical sample (designated as #1 and #2) were subjected to cyclic sublethal treatment of UV 222 or 254 followed by virus replication by zebrafish embryos to P6 independently, in comparison with passaging until P6 without treatment as control (Fig. 5). To avoid overinterpretation of results due to inherent errors in sequencing, we used a minimum of read depth of 20, and variant frequency threshold of 20%, along with two workflows to identify SNPs.

**Fig. 5.**
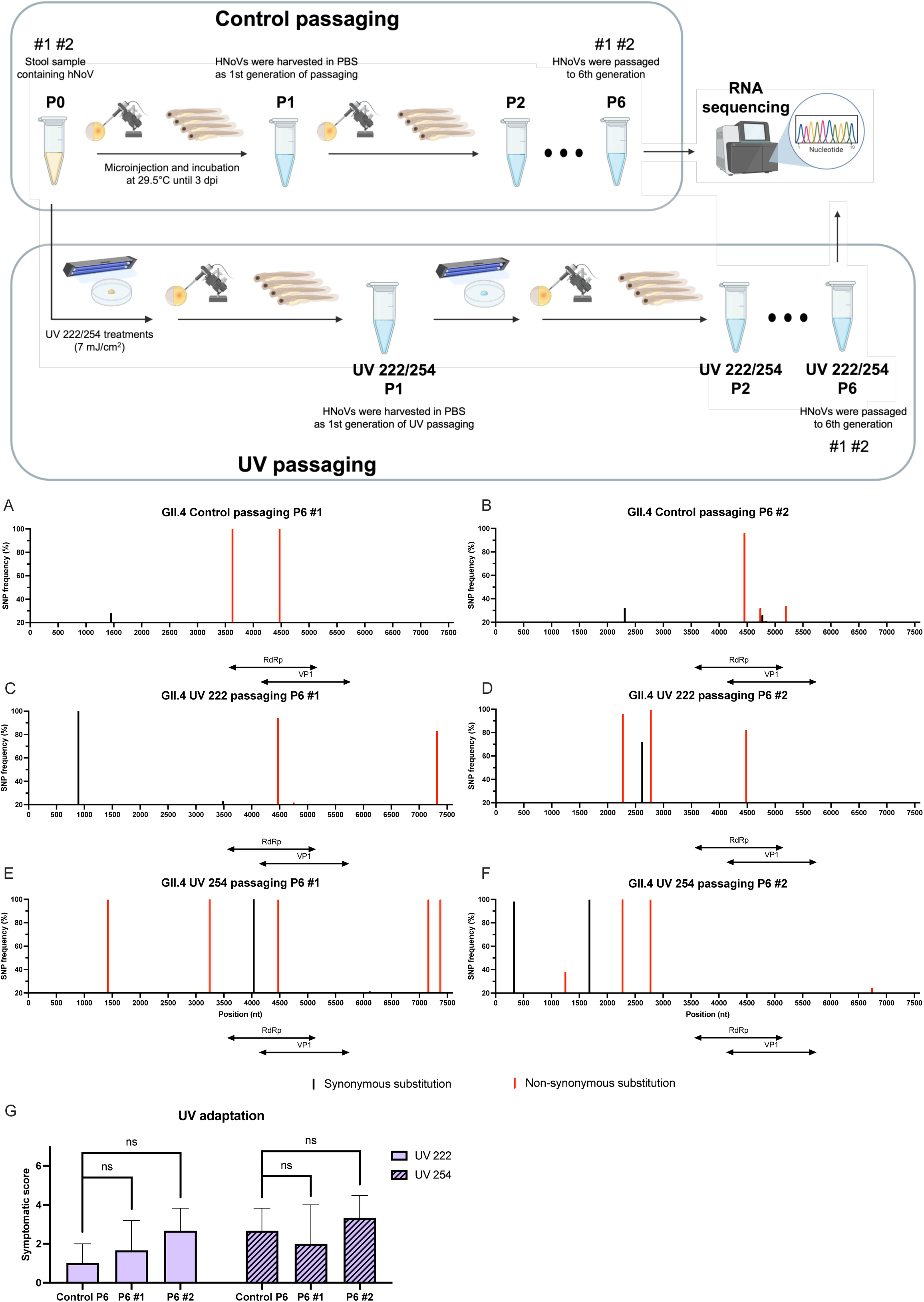
Illustration of control and UV passaging of hNoV GII.4 Sydney[P16] in zebrafish embryo model in duplicates, together with SNPs identified in 6th generation (P6) of control (A, B), UV 222 (C, D) and UV 254 (E, F) compared to stool sample (P0) in two independent passaging lines (#1 and #2), and the sum symptomatic scores of eight zebrafish larvae injected with P6 either in control and UV passaging (G), which were treated with UV 222 or 254 at dose of 7 mJ/cm^2^. Two variant determinant regions: RNA-dependent RNA polymerase (RdRp) is located within 3558–5087 nt and open reading frame 2 in 5071–6693 nt. Red columns are SNVs with non-synonymous substitution; black columns are SNVs with synonymous substitution; The threshold of SNP frequency is 20%. ns: not significantly different, *p* > 0.05.

In general, UV 222 (Fig. 5C and 5D) and UV 254 (Fig. 5E and 5F) induced more variations in hNoV GII.4 Sydney[P16] genome than control after sixth generations (Fig. 5A and 5B), especially for non-synonymous substitutions. However, in the two variant determinant regions of RNA-dependent RNA polymerase (RdRp; 3558–5087 nt) and ORF2 (5071–6693 nt), the number of SNVs didn’t increase for UV treated samples. In the control samples, two SNVs with non-synonymous substitutions were observed at 3627 and 4471 nt in sample #1, while only one SNV at 4471 in #2. All three SNVs were located within RdRp. In contrast, there appeared three non-synonymous substitutions in UVC 222 #1 (Fig. 5C), 4471 and 4753 nt in RdRp and 7323 nt in ORF3 (6693–7499 nt). Additionally, three non-synonymous substitutions were identified in UV 222 P6 #2, all from ORF1 (3–5090), namely at 2273, 2774, 4480 nt (Fig. 5D). Five non-synonymous mutations in UV 254 #1 were at 1422, 3244, 4471 nt in ORF1, together with 7159 and 7374 nt in ORF3 region (Fig. 5E). From UV 254 #2 (Fig. 5F), three SNVs with non-synonymous substitutions were located at 1250, 2273, 2774 and 6741 nt. One unique deletion was found at 320 nt (GCTGGTTCT→GCT), removing two amino acid glycine and serine in p48 region. When exposing UV P6 samples of GII.4 Sydney[P16] to both sublethal UV treatments, all of them didn’t shown difference in symptomatic scores compared with control P6, suggesting no adaptation of hNoV (Fig. 5G).

In comparison, the host environment in our study (*Supplemental material* S5) may represent a more critical driver for viral evolution, especially in inducing mutations within variant-determinant regions, including RdRp and ORF2, which respectively define the P-type and G-type classifications of hNoV (36). The numbers and frequencies of non-synonymous SNVs observed after six consecutive cycles of sublethal UV exposure and passaging were lower than those accumulated through extended viral passaging up to 24th generation. Moreover, both the numbers of SNVs and the overall mutation frequencies of hNoV detected in zebrafish in this study, were far lower than the levels reported in chronically hNoV-infected patients (37, 38). Altogether, these findings suggest that while sublethal UV exposures may contribute to limited evolutionary changes, the host environment plays a more dominant role in triggering hNoV variants.

## Discussion

To date, the absence of standardized cell lines for hNoV has continued to limit the direct evaluation of viral infectivity during disinfections. In this study, we successfully established symptom-based method for quantifying infectivity loss of four hNoV strains through the zebrafish embryo/larvae tool combined with a microinjection procedure. This approach demonstrated a capacity of quantifying up to ∼2-log reductions of hNoV inactivation, identical to the method based on the measurement of viral loads (21). Compared with the previous method based on viral loads, the new method based on symptom scores improved the linear correlation between symptom scores and initial viral loads, thereby enabling more precise quantification within the 2-log reduction range. In addition, the scoring method, which is primarily based on visual observation, eliminates the labor-intensive and costly steps involved in RNA extraction and RT-qPCR quantification. To minimize subjectivity and observational bias, clear scoring criteria were developed based on the extent of morphological abnormalities in the fish, as shown in Fig. 1. A single-blinded setup is highly recommended for the scoring process. With this scoring approach, we compared the efficacy of UV 254 and UV 222 at 7 mJ/cm^2^ and 70 mJ/cm^2^ respectively in inactivating hNoVs. Notably, the incident fluences of UV 254 and UV 222 were determined by a chemical actinometry using potassium iodide, which was applicable to UVC irradiation range (39). However, discrepancies of absolute values might occur when compared to the readings of radiometers (40).

The hNoVs tested in this study showed strain-specific variability in response to UV treatments. Overall, GII.2[P16] was considered as the most susceptible strain, while GII.17[P31] was the least susceptible one against both UV 254 and UV 222 within the four tested hNoV strains. Such variabilities within hNoV strains have also been reported previously under other inactivation conditions. For instance, GII.2[P16] showed greater thermal resistance to 95 °C for 5 min than GII.4 Sydney[P16] through RNase-RT-qPCR detection (22). Regarding chlorine disinfection, genotypes of GI.1, GII.4 New Orleans, and GII.4 Sydney demonstrated lower susceptible to NaClO than other strains like GII.6 and GII.13 as determined by RNase-RT-qPCR (15). Since the mid-1990s, GII.4 has become the predominant genotype globally, accounting for approximately 70–80% of hNoV infection cases (41). A few non-GII.4 genotypes, however, have also emerged in recent years. The recombinant strain, GII.2[P16], has been implicated in widespread outbreaks across China, Japan, and Germany (42). GII.3[P12] has been frequently associated with sporadic cases and foodborne outbreaks in both developed and developing countries (43). Notably, GII.17[P31], identified in this study as the most resistant to UVC treatment, emerged as a major cause of gastroenteritis outbreaks in China and Japan during the winter of 2014–2015, and was subsequently detected in the United States, Europe, and Australia (44). Recent surveillance indicates that GII.17 has become a predominant strain in certain regions, even surpassing GII.4 in prevalence (45). Its enhanced resistance to UVC irradiation may be one of the contributing factors to its widespread circulation and persistence.

When the viruses were tested in droplets deposited in Petri dishes, UV 222 showed comparable, if not better, efficacy in reducing hNoV infectivity and RNA integrity, and is considerably more efficient in damaging viral capsid protein than UV 254. Likely, the overlapping action of both wavelengths on viral RNA genome explains why no synergistic effect was observed in our study when the viruses were first exposed to UV 222 followed immediately by UV 254 (*Supplemental material* S4). It is known that virus inactivation at longer UVC wavelengths correlates with RNA absorption spectrum, (46, 47); whereas protein absorption becomes the dominant mechanism in the shorter wavelength region (below 240 nm) (48, 49). Thus, UV 222 in our study could primarily destroy viral capsid proteins that strongly absorb UV photons at the wavelengths, transferring energy or inducing secondary damage to viral RNA genome, ultimately resulting in infectivity loss (50). With approximately 14% higher photon energy than UV 254 as reported previously, UV 222 has a greater potential to break chemical bonds within capsid proteins (7, 51). This is supported by enhanced absorption of amino acid (e.g. histidine) and photodegradation of oligopeptides (e.g. angiotensin II) under 222 nm irradiation (48). Similar observations have been reported for MS2 as an hNoV surrogate (16, 52) and were consistently confirmed in this study. However surprisngly, TV, a much more commonly used surrogate virus for hNoV in inactivation studies in recent years (1, 22, 53), demonstrated opposite trends and showed exceptional resistance to UV 222. This unique property of TV may reflect compositional and structural differences in its capsid proteins, leading to reduced far-UVC absorbance and limited energy transfer to the viral RNA (50). It has been reported previously that the peptide bonds in protein backbone are related to their differential absorbance in far-UV region (54). These results underscore the necessity of evaluating genuine hNoV strains to ensure the reliability of inactivation strategies.

When the viruses were spiked onto stainless steel surfaces and porcine skin, UV 222 showed comparable inactivation efficacy to UV 254 on hydrated virus inocula. When the viruses were dried on the surfaces or mixed with simulated vomitus, the inactivation efficacy of both UV 222 and UV 254 was markedly reduced, with UV 254 proving more effective than UV 222. We assume this was mainly due to the low penetration of UVC, which represents a key limitation of this technology for environmental disinfection. Compared with Petri dishes, stainless steel has a rougher surface, while porcine skin possesses a biological structure that provides indentations, allowing dried viruses to remain partially shielded from direct UV exposure. This result aligned with prior report testing UV 222 on dried human rhinovirus and human coronavirus on glass carriers (55). The organic matters in the simulated vomitus could also have physically shielded the viruses from UV exposure, being consistent with report on SARS-CoV-2 tested in human saliva (47). In addition, it has been reported that reactive oxygen species (ROS) are generated upon UV 222 treatment (56), which may provide an additional explanation for why UV 222 outperformed UV 254 in inactivating hydrated hNoV in this study.

UV 222 presents a safer exposure profile for human skin and eyes (8, 26). Lately, a new study on the safety profile of far-UVC 222 agreed with previous reports that minor DNA damage at high doses (around 500 mJ/cm^2^) was induced evaluated using live human skin biopsies (8, 57). Also, well-filtered UV 222 lamp produces very low levels of ozone (less than 10 ppb) in real-world scenarios (58). These unique features make it a promising alternative for replacing conventional UVC light for surface disinfection regarding hNoV control. The installation of 222 nm UV lights in enclosed environments such as cruise ships, food processing plants, and healthcare facilities can enable real-time, continuous disinfection even during occupancy (59, 60). Moreover, its demonstrated safety property supports direct applications into high-touch surfaces or even for hand disinfection. However, this study identified a trade-off between safety and efficacy. While UV 222 is safer for human exposure than UV 254, its lower penetration makes it more susceptible to interference from surface roughness and organic matter, reducing its inactivation performance under certain environmental conditions. The shadow effect may hinder uniform exposure, reducing antiviral efficacy on irregular or shaded surfaces (61). Furthermore, hNoV is frequently transmitted via fecal or vomitus droplets containing complex organic matrices, such as proteins and carbohydrates, which can absorb or scatter UV photons, thereby protecting virus from irradiation (62). Similar attenuation effect have been observed in water treatment system with elevated turbidity or suspended solids (63). Therefore, UV 222 should be applied after surface cleaning to remove the organic soils, and ideally on moist surfaces to maximize its efficacy. Moving forward, standardizing far-UVC device performance, dose calibration, and safety criteria will be critical for ensuring consistent and effective field implementation. Moreover, combining UV 222 with complementary disinfection methods such as chemical sanitizers, photocatalytic coatings, or aerosol treatments less affected by shadowing, may further enhance its reliability and broaden its applicability in real-world settings.

## Materials and methods

### The propagation and titration of surrogate viruses

TV stock was kindly provided by Professor Xi Jiang at Cincinnati Children’s Hospital Medical Center (Cincinnati, OH, USA). The propagation and titration of TV were performed using the monkey kidney cell line LLC-MK2 (ATCC^®^ CCL-7™). Cells were cultured in M199 medium (Gibco™ 11150, Life Technologies Holdings Pte Ltd., USA) supplemented with 10% fetal bovine serum (qualified, heat inactivated, Gibco™) and 1% penicillin streptomycin (Gibco™). Cultures were maintained at 37°C in a 5% CO_2_ incubator (CelCulture^®^ CCL-170/240L, ESCO Micro Pte. Ltd., Singapore). Virus titer were determined using the 50% tissue culture infectious dose (TCID_50_) method (53). For serial passaging from sixth to fifteenth generation (P6-P15), confluent monolayers of LLC-MK2 cells were infected with TV at a multiplicity of infection (MOI) of 0.05 and incubated at 37°C in atmosphere aspirated with 5% CO_2_ for 3 d until 80% cytopathic effect (CPE). The infected cells were subsequently subject to three freeze-thaw cycles and centrifuged at 8000 *×g* for 10 min at 4°C. The resulting supernatant was titrated and stored at −80°C for future use.

*Escherichia coli* 15579 and MS2 coliphage (*E. coli* bacteriophage 15597-B1™) were both obtained from the American Type Culture Collection (ATCC; Rockville, MD, USA). The titer of MS2 phage was examined by a plaque assay method (53). Briefly, 100 μL of virus solution was plated onto double-layer tryptone yeast glucose agar (TYGA; Oxoid, Thermo Fisher Scientific Inc., Basingstoke, UK), with *E. coli* incorporated in the semisolid overlayer. Plaques were enumerated and expressed as plaque-forming units per mL (PFU/mL). For passaging, 10 mL freshly cultured *E. coli* was infected with MS2 to achieve a final concentration of 1.0×10^7^ PFU/mL. Cultures were incubated at 37°C with shaking at 150 rpm for 24 h and were subsequently centrifuged at 8000 ×*g* for 10 min at 4°C. The resulting supernatant was titrated and stored at −80°C for future use.

### HNoV titration and analysis

#### HNoVs from stool samples

Stool samples containing hNoV genotypes GII.2[P16], GII.4 Sydney[P16] and GII.17[P31](Pe) were kindly provided by the Molecular Laboratory, Department of Molecular Pathology, Singapore General Hospital. Ten percent GII.3[P12] stool suspension was obtained from Institute of Agrochemistry and Food Technology (IATA-CSIC), Spain. Following the protocol described by H. Tan Malcolm Turk et al. (21), hNoV samples were prepared for microinjection by suspending 100 mg of stool sample in 1 mL sterile phosphate-buffered saline (PBS; Vivantis Technologies Sdn. Bhd., Selangor, Malaysia). The suspension was vortexed thoroughly and centrifuged at 9000 ×*g* for 5 min. The resulting supernatant was titrated by RT-qPCR and stored at −80°C for future use.

### Zebrafish husbandry, embryo maintenance and microinjection in *Supplemental material* S1

#### Microscopic observation

At 2 dpi, eight zebrafish larvae injected with hNoVs were individually transferred into the wells of 96-well plate. At 3 dpi, larvae were anaesthetized for 2-3 min using 0.2 mg/mL Tricaine (MS-222; Sigma-Aldrich, St. Louis, MO, USA) prepared in E3 buffer to facilitate the observation of the yolk sac and cardiac region. Morphological and behavioral characteristics, including abnormality in the yolk sac area, head and pericardial region, the sign of edema, tissue damage, along with responsiveness to stimulation, were subsequently recorded (64, 65). A stereo microscope (Nikon SMZ25, Nikon Instruments Inc., Tokyo, Japan) equipped with a digital camera (Nikon DS-10) was used for downstream analysis.

#### Virus harvest

Zebrafish embryos or larvae injected with hNoV stool samples were harvested either in pools of ten individuals per sample for passaging and titration, or individually for validation of viral loads per fish. Following euthanasia on ice, the harvested fish samples were transferred into 1.5-mL microcentrifuge tubes containing 300 μL PBS, and homogenized using a FastPrep^®^-24 5G tissue and cell homogenizer (MP Biomedicals, Irvine, CA, USA). The homogenization protocol consisted of three cycles of 15 s at 6500 rpm, with 60 s intervals. Homogenates were subsequently centrifuged at 8000 ×*g* for 10 min, and the resulting supernatants were collected as the first generation of hNoV passaging (P1). Further passaging was conducted using the same protocol. All hNoV samples harvested from zebrafish were stored at −80°C for further analysis (RNA extraction, and UV inactivation studies). HNoV P3 to P6 were applied in the inactivation study.

#### RNA extraction

Total RNA isolation was performed using the RNeasy Mini Kit (Qiagen, Hilden, Germany), following the manufacturer’s protocol.

#### RT-qPCR

As the gold standard for hNoV detection (66), RT-qPCR was employed in this study to assess the effects of the UV treatments and to compare performance with other detection methods. RT-qPCR was performed using GoTaq^®^ Probe 1-step RT-qPCR system (Promega, Madison, WI, USA). The primers and probes used for the three tested viruses are listed in Table 1. Cycling conditions were as follows: 45°C for 15 min, 95°C for 10 min, followed by 40 amplification cycles of 95°C for 15 s and 60°C for 30 s. Cycle threshold (C_T_) values were determined using StepOnePlus™ Real-Time PCR System (Applied Biosystems, Foster City, CA, USA), reflecting the viral dose level (21). For standard curve construction, please refer to *Supplemental material* S1.

**Table 1.**
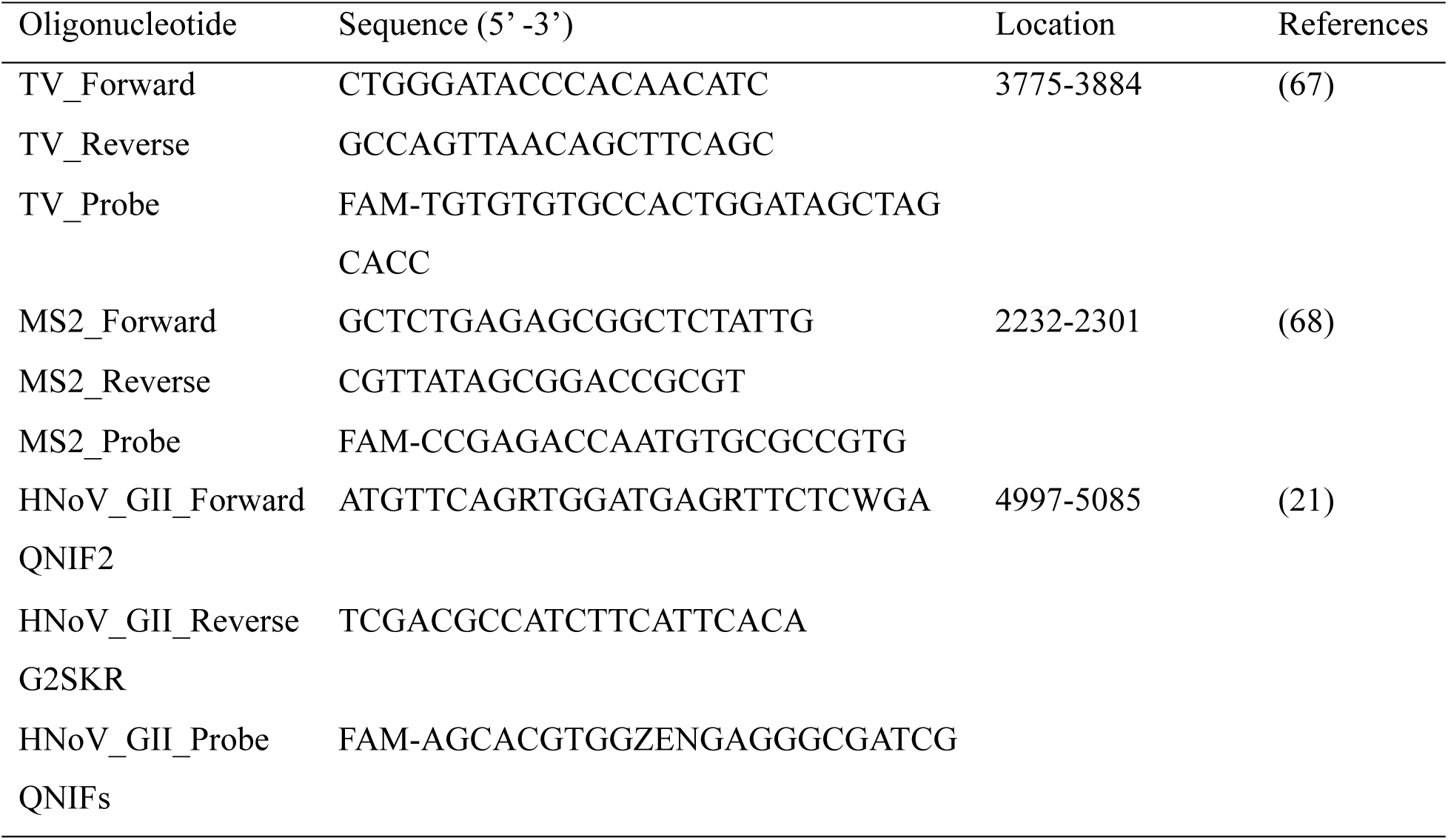
Primers and probes for TV, MS2 and hNoV.

#### LR-RT-qPCR

To assess the RNA genome integrity following UV treatments, LR-RT-qPCR enables the synthesis and quantification of longer complementary DNA (cDNA) from intact RNA, while fragmented RNA prevents amplification. Long-range reverse transcription was performed using SuperScript™ III Reverse Transcriptase (Invitrogen, Thermo Fisher Scientific, Waltham, MA, USA), adapted from the manufacturer’s instruction and D. Li et al. (32). Each 20-μL reaction volume contained 200 U reverse transcriptase, 40 U Recombinant RNasin® Ribonuclease Inhibitor (Promega), 0.5 mM deoxynucleotide triphosphates (dNTPs; Promega), 500 ng oligo(dT)_15_ Primer for hNoVs and TV (Promega), and reverse primer for MS2 (5’-AATCCCGGGTCCTCTCTTTA-3’, location 3509-3528) (69). First-strand cDNA synthesis was initiated by mixing RNA sample with primers and dNTPs, followed by incubation at 65°C for 10 min and rapid cooling at 4°C for at least 1 min using a thermal cycler (Bio-Rad T100™, Hercules, CA, USA). The remaining reagents were then added, and reverse transcription was carried out at 50°C for 60 min, followed by enzyme inactivation at 70°C for 15 min. To remove cRNA, RNase H (2 U/μL; Invitrogen) was added and incubated at 37°C for 20 min. Finally, GoTaq^®^ Probe RT-qPCR system was used to quantify the cDNA.

#### RNase-RT-qPCR

To assess capsid integrity following UV treatment, RNase digestion was carried out prior to RT-qPCR analysis to selectively remove viral RNA that was either fully exposed or enclosed within damaged capsids. Specifically, 40 units of RNase ONE™ ribonuclease (Promega) were added to the treated virus samples (70), followed by incubation at 37°C for 20 min. After digestion, the reaction was terminated by the addition of lysis buffer in RNA extraction.

### UV treatments

#### General set-up of UV treatment

UV treatments were performed in a Class II biosafety cabinet (LA2-4A1, ESCO Micro Pte. Ltd., Singapore) with a low-pressure mercury lamp of 254 nm UV radiation (UV-30) for UV 254 treatment. For 222 nm radiation, a KrCl excimer lamp (ST28-15W-222, Shenzhen Suntech Co., Ltd., China) was employed. Measurement of incident fluence rate of UV irradiations along with emission spectra of UV lamps are provided in *Supplemental material* S3.

To ensure comparable exposure conditions, the distance of UV 222 lamp to treatment area was adjusted, so that the incident fluence rates of both wavelengths were equivalent. Subsequently, 10 μL droplets of virus samples were deposited in Petri dish and exposed to UV irradiation at a sublethal dose of 7 mJ/cm^2^ (> 90% inactivation of MS2 based on our preliminary data) or at a virucidal dose of 70 mJ/cm^2^ (> 99.99% inactivation of MS2 based on our preliminary data). Treatments were applied using either UV 222 or UV 254 individually. After UV treatments, the droplets were recollected by pipetting into 1.5-mL tubes for RNA extraction or infectivity assays. The virus recovery rates evaluated by RT-qPCR were between 96.8% to 99.3% (Table S1).

#### Simulation of surface disinfection scenarios

Stainless steel disks were purchased from Heshengan Stainless steel Co., Ltd.. Fresh porcine ears were purchased from local food market with the hair removed in advance. The central outside portions of the ear were washed with PBS, soaked in 70% ethanol for 30 min and cut into 1 cm × 1 cm pieces. Subsequently, the porcine ear skin pieces were subject to UV treatment for 30 min in the biosafety cabinet before use. Simulated vomitus was prepared according to the protocol of INFOGEST 2.0 *in vitro* digestion up to gastric phase (71) using 100 mL milk and 10 g instant oats as the substrate. After the simulated digestion, the mixture (pH = 5.4) was centrifuged at 9000 ×*g* for 10 min and subsequently filtered through 10 μm membrane filters.

MS2 or hNoV GII.17[P31] were diluted in deionized water or simulated vomitus to achieve approximately 10^8^ PFU/mL or genomic copies/mL, respectively, and 10 μL virus solution was deposited onto a stainless steel disk or porcine ear skin piece as one sample. As hydrated samples, the droplets were directly exposed to 70 mJ/cm^2^ UV 222 or UV 254. To recover the viruses, the treated droplets were transferred into 1.5 mL microcentrifuge tubes using a pipette. The recovery rates were evaluated by RT-qPCR for hNoV GII.17[P31] and by plaque assay for MS2, and were consistently above 97% as shown in Table S1. As dried samples, the droplets were allowed to dry for 2 h in biosafety cabinet, followed by the UV treatments. To recover the viruses, 30 μL of sterile deionized water was pipetted onto the dried, virus-spiked spot, aspirated and dispensed five times, and then transferred into 1.5 mL microcentrifuge tubes. The recovery rates of dried viruses were around 50% from stainless steel discs, and above 5% from porcine ear skin.

### Evaluation of adaptation and mutation rates after repeated UV treatments

To address the concern regarding UV adaptation in hNoV and its potential role in driving viral evolution, the repeated exposures to sublethal doses of UV irradiations along passaging were conducted, followed by the evaluation of inactivation rates and variant calling analysis. The strain GII.4 Sydney[P16] was employed due to the global epidemiological dominance and high evolutionary rates (72). Specifically, droplets containing hNoV GII.4 Sydney[P16] were first subjected to UV 222 or 254 (7 mJ/cm^2^), and subsequently injected into zebrafish embryos for passaging. This procedure was repeated until sixth generation (P6) within two independent lines as UV passaging samples (Fig. 5). In parallel, untreated samples were serially passaged to P24 as control for comparative analysis (Fig. S3). Afterwards, UV P6 samples exposed to UV 222 or 254 to were evaluated using symptomatic scoring to assess the inactivation rates of hNoV in comparison to control P6 subject to the same treatments.

Referring to the previous studies on single nucleotide polymorphisms (SNPs) and viral evolution analysis (30, 73), the sequencing and bioinformatics analysis were as follows. The virus samples were processed with RNA extraction steps and sequenced though Illumina NovaSeq 6000 platform with paired end (150 bp). The obtained NGS raw data following quality control was approximately 20 million reads (6 GB), which were deposited in the NCBI Sequence Read Archive (SRA) under accession number [PRJNA1256631]. Host (zebrafish) RNAs were removed using Bowtie2 to avoid contamination and potential misinterpretation of mutations. Raw reads from P0 of GII.4 Sydney[P16] sample (stool sample) were uploaded onto Galaxy (Version: 24.2.1.dev0) and assembled with MEGAHIT. The assembled contig of 7566 kb was deposited as a BLAST query in NCBI databases, indicating 98.37% similar to the complete genome of hNoV GII (NC_039477), which was set as reference genome. Subsequently, two distinct workflows of alignment and variant calling were concurrently performed. Firstly, raw reads from control or UV treated samples (GII.4 Sydney[P16] P6 as control, UV treated P6 samples as treatment groups) were aligned to reference genome with Bowtie2, and variant calling was performed using VarScan. The filtration parameters were as follows: minimum read depth 10, minimum supporting reads 5, minimum base quality at a position to count a read 25, minimum variant allele frequency threshold 20%, *p*-value threshold for calling variants 0.05, others as default. Secondly, BWA-MEM2 was used for alignment, followed by lofreq for variant calling. The minimal coverage was 10, minimum base quality or quality for alternate bases 25, minimum mapping quality 25, minor variant frequency threshold 20%, others as default. Last, only those single nucleotide variants (SNVs) and insertions and deletions (indels) identified by both workflows were further analyzed and annotated with SnpEff (30, 73, 74). For variant calling analysis agreed by VarScan and lofreq, each sample achieved an average sequencing depth exceeding 100×, notably higher than the recommended depth of 50× (75), ensuring high-confidence variant detection.

### Statistical analysis

The effects of 222 or 254 nm UVC lights across hNoV strains or surrogates, as well as the application on surfaces were analyzed through two-way analysis of variance (ANOVA) and Tukey’s HSD test in RStudio (Version: 4.4.2). The differences among the 222 nm groups were denoted using lowercase letters, whereas those in the 254 nm groups were capitalized. For UV adaptation, one-way ANOVA was used in each treatment. The differences were denoted with asterisks. All figures were generated in GraphPad Prism (Version: 10.4.1).

## Acknowledgement

This research was financed by a department fund at the Department of Food Science and Technology, National University of Singapore (E-160-00-0015-01, Food Microbial Safety Research, Dr. Dan Li) and the Biomedical and Health Technology Platform, National University of Singapore (Suzhou) Research Institute (Dr. Dan Li). The authors would like to thank Dr. Zhihui Yang from Division of Molecular Biology, Center for Food Safety and Applied Nutrition U.S. Food and Drug Administration for her generous and insightful technical suggestions.

## Supplemental Materials

Supplementary methods and results.docx

